# Isolation and characterization of bacteriophages that infect *Citrobacter rodentium*, a model pathogen for investigating human intestinal diseases

**DOI:** 10.1101/248153

**Authors:** Carolina M. Mizuno, Tiffany Luong, Robert Cedarstrom, Mart Krupovic, Laurent Debarbieux, Dwayne R. Roach

## Abstract

Enteropathogenic *Escherichia coli* (EPEC) is a major etiology for diarrheal diseases among children. Antibiotics, when used appropriately, are effective; however, their overuse and misuse has led to the rise of antibiotic resistance worldwide. Thus, there are renewed efforts into the development of phage therapy. Due to the drawbacks of EPEC *in vivo* models, a surrogate is the mouse-restricted gut pathgoen *Citrobacter rodentium*. In this study, two new phages CrRp3 and CrRp10, which infect *C. rodentium*, were isolated and characterized. CrRp3 was found to be a new species within the genus *Vectrevirus* and CrRp10 is a new strain within the genus *Tequatrovirus.* Neither phage carries known genes associated with bacterial virulence, antibiotic resistance, or lysogeny. CrRp3 and CrRp10 appear to have independently evolved from *E. coli* phages. CrRp3 appears to be the more ‘potent’ being 24x more likely to find a host cell and has a shorter lytic cycle, while CrRp10 at MOI 0.001 was able to maintain bacterial density below the limit of detection after 18 h. We found that hypoxia (5% O_2_ and 5% CO_2_) inhibited CrRp3 ability to reverse exponential bacterial growth. It is unclear whether the subtle characteristic differences between CrRp3 and CrRp10 will influence treatment efficacy in future phage therapy *in vivo* investigations.

## INTRODUCTION

Diarrheal diseases continue to be one of the foremost public health issues globally, responsible for more than 1.6 million deaths each year [1]. While mortality rates have reduced substantially (by 34% among children <5 years and by 21% among all people) over the last two decades, the incidence of diarrhea has not reduced nearly as much (10% reduction in children and 6% reduction overall) [1]. Enteropathogenic *Escherichia coli* (EPEC) is a major etiology for diarrhea morbidity and mortality among children <5 years [1,2]. When used appropriately, antibiotics can reduce the severity and duration of diarrheal diseases; however, overuse and misuse of antibiotics in the treatment of diarrhea has led to an alarming increase in antibiotic resistance (AMR) globally [1-3]. These observations illustrate that diarrhea mortality is largely avoidable and renewed efforts to reduce disease burden with non-antibiotic strategies is urgently needed.

Mouse models for study of EPEC diseases have had limited utility because gastrointestinal microbiota generally defends well against human *E. coli* infection [4]. Consequently, most mouse models of *E. coli* gastrointestinal diseases require substantial antibiotic pre-treatment to reduce resident microbiota [5-8]. The preferred surrogate for EPEC diseases is the mouse-restricted pathogen *Citrobacter rodentium* [9]. Both EPEC and *C. rodentium* cause attaching and effacing (A/E) lesion and share the same pool of locus of enterocyte effacement (LEE)-encoded and non-LEE-encoded effector proteins to subvert and modulate gut epithelial barrier properties [9,10]. Importantly, without the aid of antibiotics, *C. rodentium* caused pathologies indistinguishable from those observed in the human EPEC gut infection [11].

Bacteriophages (phages) are one of the foremost antibacterial agents under development and clinical testing for treating life-threatening AMR pathogens [12,13], with several recent successful compassionate use phage therapy cases (For examples see [14-16]). Phages are highly abundant and ubiquitous viruses that only infect and kill bacteria. Other promising features of phages include a narrow host range, self-replication at sites of infection, disruptive biofilm activity, and potential synergy with classical antibiotics [13,17,18]. Traditionally, virulent double-stranded DNA tailed phages that carry out a strictly lytic infection cycle are considered the ideal viral type for human therapeutic applications [12,13]. However, only four virulent phages that infect *C. rodentium* have been described to date, including phiCR1 [19], R18C [20], CR8 and CR44b [21]. In addition, *in vivo C. rodentium* phage therapy has not been explored.

In this study, we characterize two new virulent phages CrRp3 and CrRp10 that infect *C. rodentium*. Using genome sequence annotations, phenotypic susceptibility under normoxic and hypoxic environments, host cross-genera infection ranges, and phylogenetic analysis, we assess their subtle feature differences that may improve efficacy in future therapeutic evaluation in a A/E diarrheal disease mouse model.

## MATERIALS AND METHODS

### Strains and culturing

*C. rodentium* strain ICC180 [22] was grown in Luria-Bertani (LB) broth under either normoxic (atmospheric 21% O_2_ and 0.04% CO_2_) or hypoxic (5% O_2_ and 5% CO_2_) environmental chambers at 37°C with orbital shaking, as needed. Prior to hypoxia culturing, LB was precondition at 5% O_2_ and 5% CO_2_ for 2 days to remove dissolved oxygen in the medium. Agar (1.5%) was added to LB for solid growth medium.

### Phage isolation, cultivation and purification

*C. rodentium* phages were isolated from 0.25 μm syringe filter (Sartorius) sewage sample collected from water treatment plants around Paris France. Individual plaque forming units were isolated from a 10 μL sample spread over LB agar Petri plate, overlaid with 2 ml of mid-log growing ICC180 under normoxic conditions. After overnight incubation, a sterile Pasteur pipette was used to select visible PFU with different morphologies and phages enriched by culturing with ICC180 in fresh LB. PFU selection and enrichment was repeated 5 times. Final phage lysates were sterilized by high-speed centrifugation and 0.2 µm filtration. For electron microscopy, phage lysates were further purified by cross-flow filtration (CFF) and cesium chloride (CsCl) density gradient ultracentrifugation using a methodology previously described [23].

### Transmission electron microscopy

To visualize viral morphology purified phages in TN (10 mM Tris-Cl, 10 mM NaCl) buffer were placed onto carbon-coated copper grids and negatively stained with 2% (wt/vol) uranyl acetate for 30 s. Then, phages were visualized using a 120 FEI-1 Tecnai Biotwin TEM at an accelerating voltage of 120 kV and a Gatan Orius 1000 digital camera recorded the micrographs.

### DNA extraction, and nucleotide sequencing

Genomic DNA was extracted from sterile DNase and RNase pretreated purified phages by a phenol-chloroform extraction as previously described [24]. Sequencing libraries with single index were prepared using the NEBNext DNA library prep kit (New England BioLabs, Ipswich, MA) and then sequenced on the Illumina MiSeq sequencing platform (Illumina, San Diego, CA, United States) with paired-end 300 nucleotide reads. Raw reads were trimmed by FastQC v10.1 (www.bioinformatics.babraham.ac.uk/projects/fastqc/) and *de novo* assembled using the CLC Assembler (Galaxy Version 4.4.2).

### Nucleotide alignments, amino acid alignments, annotations and phylogenetic analyses

Annotation of the specific function of ORFs was conducted using rapid annotations of subsystems technology (RAST) [25]. The presence of transfer RNA (tRNA)-encoding genes was determined using the tRNAscan-SE database [26] and protein-coding genes were predicted using Prodigal [27]. Additional annotation of genes were done manually on the HHPRED server [28] and by comparing against the NCBI NR, COG [29], and TIGRfam [30] databases. Comparative analysis among related complete genomes were performed using tBLASTx or BLASTN [31]. Phylogenetic relationships of phages CrRp3 and CrRp10 were determined using VICTOR [32] and pairwise comparisons of the nucleotide sequences were conducted using the Genome-BLAST Distance Phylogeny (GBDP) method [33] under settings recommended for prokaryotic viruses [32]. GBDP phylogenetic relationships were constructed from *Citrobacter, Escherichia* and *Synechococcus* podoviruses and myoviruses downloaded from GenBank (12/21/2017; Supplementary Table S4). The resulting intergenomic distances (100 replicates each) were used to infer a balanced minimum evolution tree with branch support via FASTME including SPR post processing [34]. The trees were rooted with *Synechococcus* phages (outgroup) and visualized with FigTree V1.4.3 (http://tree.bio.ed.ac.uk).

### Phage adsorption

A similar protocol [35] was adopted to determine phage adsorption rate. In 9 ml of LB, 2.5×10^8^ colony forming units (CFU) mL^-1^ of exponentially grown ICC180 was mixed with 1 mL of phages at a multiplicity of infection (MOI) of 0.001 and incubated at 37° with constant shaking. Every minute supernatant samples were withdrawn, serially diluted, and titrated. Phage adsorption rate was estimated by the equation k=2.3/Bt*log(*Po*/*P*), where k is the adsorption rate constant, in phage^-1^ cell^-1^ ml^-1^ min^-1^; B is the concentration of bacterial cells; t is the time interval in which the titer falls from Po (original) to P (50%).

### One-step growth curves

We performed a one-step growth curve procedure as described previously [36], with some modifications. Briefly, exponentially grown *C. rodentium* in 9 mL LB at 2.5 ×10^8^ CFU mL^-1^ was mixed with 1 mL of phages at a multiplicity of infection (MOI) of 0.01. At regular intervals, two samples were taken every 5 min, with one used to enumerate free phages in solution; the second was treated with 1% chloroform to release intracellular phage. All samples were titrated for phage numbers. Comparison between chloroform-treated and non-treated samples showed the eclipse/latent periods and burst size.

### Phage lysis curves

*C. rodentium* strain ICC180 from overnight cultures was used to spike fresh LB and grown 37°C with shaking until an optical density (OD_600_) of ∼5. For hypoxia studies, ICC180 was grown in preconditioned LB at 5% O_2_ and 5% CO_2_. Cells were then washed twice, by centrifugation at 6,000 g for 10 min and resuspension of the pellet in chilled H_2_0. Microplate wells were filled with 100 µl of 2 times LB concentrate (preconditioned if required) before adding washed cells to give a total of 2 ×10^6^ CFU. Phages were added at different MOIs, H2O added for a total volume of 200 µL, and microplates immediately placed in a microplate reader (Clariostar; BMG) and OD_600_ measured every 6 min for 18 h. For hypoxia, O_2_ and CO_2_ levels were regulated within the platereader with an atmospheric control unit (BMG) supplied with CO_2_ and N_2_ gases.

### Phage host range

We evaluated host range using a spot test with 10^7^ PFU of phages onto dried lawns of 33 bacterial strains (listed in Table 3) that were grown to OD_600_ 0.22 on agar and then incubated at 37°C overnight.

Bacterial species tested included 27 *Escherichia coli* strains, *Erwinia carotovora* CFBL2141 [37], *Pseudomonas aeruginosa* PAK [38], *Rouxiella chamberiensis* nov [39], *Serratia marcescens* DB11 [40] and SM365 [41]. For comparison, the host ranges were also determined for *E. coli* phages LF82_P10 [42], LM33_P1 [43], AL505_P2 [44], CLB_P2 [45], 536_P7 [46] and *Pseudomonas aeruginosa* phage PAK_P1 [23].

### Statistics

Data is shown as means ± standard deviation (SD). Statistical analyses were performed using GraphPad Prism (v5). A Student’s t-test or one-way analysis of variance with Tukey’s multiple-comparison test was used to determine differences between two or multiple groups, respectively. A p <0.05 was considered significant.

### Accession numbers

The complete genome sequences of vB_CroP_CrRp3 (CrRp3) and vB_CroM_CrRp10 (CrRp10) are deposited in NCBI GenBank with accession numbers MG775042 and MG775043, respectively.

## RESULTS

### Isolation, morphology, sequencing, and genome annotation of CrRp3

The novel phage CrRp3 (formal name: vB_CroP_CrRp3), which infects *C. rodentium* strain ICC180, was isolated from sewage. Figure 1a shows that CrRp3 virion has a typical podovirus morphology, with an isometric head, likely icosahedral, and a short tail. Plaques produced by CrRp3 on ICC180 were of 2.36 ± 0.46 mm^2^, mean ± 95% confidence interval (Fig. 1b).

**Figure 1.**
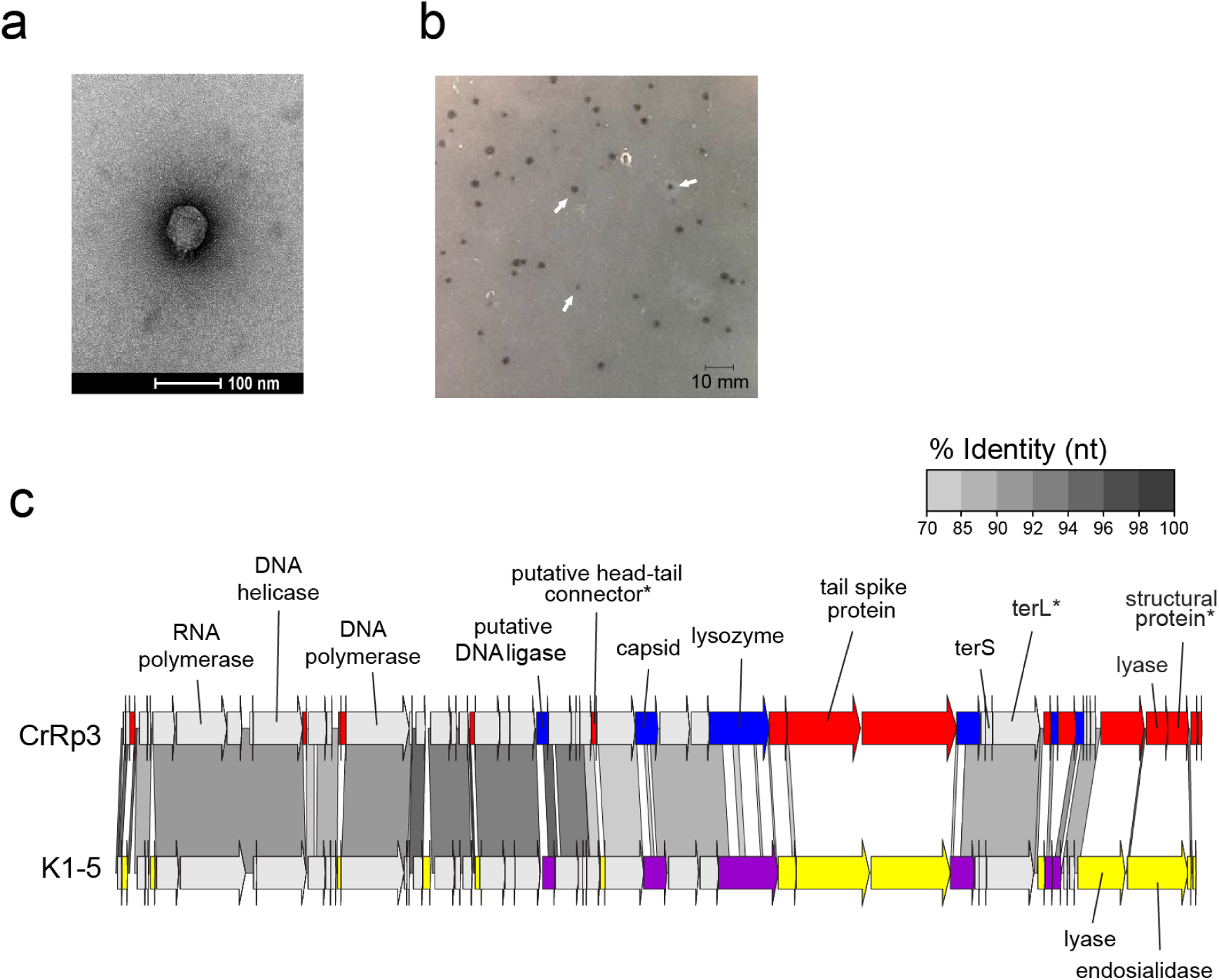
Morphology, genome organization and gene functional comparison of CrRp3. **a)** Electron micrographs of CrRp3 negatively stained uranyl acetate. **b)** Plaque forming units by CrRp3 on lawns of *C. rodentium* ICC180. **c)** Alignment of CrRp3 and closest taxonomic relative *E. coli* phage K1-5. Genes are colored according to sequence homology, with genes exclusive to CrRp3 labeled red, genes exclusive to K1-5 labeled yellow, and genes that are homologous between CrRp3 and K1-5 but highly variable labeled blue.

Next generation sequencing showed that CrRp3 has a terminally repetitive dsDNA genome of 44.3 kb with 54 predicted coding sequences (CDSs) (Fig. 1c and Table 1). However, only 19 (35%) could be assigned a putative function (Table S1). Interestingly, CrRp3 has a GC content of 45%, which is 10% lower than in its host *C. rodentium*, which has a GC content of 54.5%. The genome does not encode proteins with identifiable sequence homology to known lysogeny-associated proteins, suggesting that CrRp3 is virulent with a strictly lytic lifecycle. In addition, CrRp3 genome does not carry recognizable virulence-associated genes or antibiotic resistance genes.

**Table 1.**
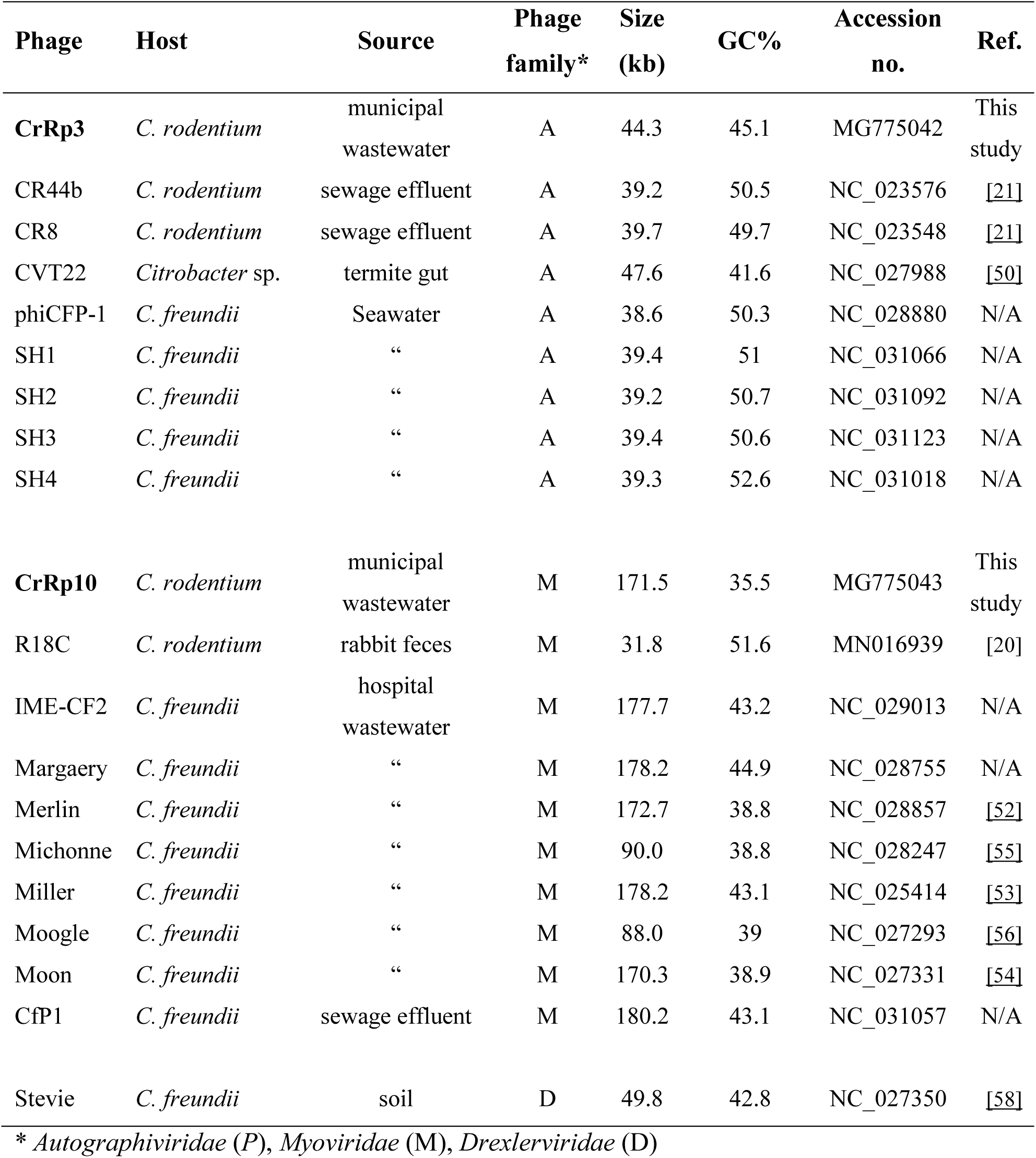
Citrobacter phages and their genome features.

At the time of analysis, the closest taxonomic relative of CrRp3 is the *E. coli* Sp6-like phage K1-5. The two phages share both gene content and genome organization with 90% (over 76% of the genome) and 77% (over 75% of the genome) pairwise nucleotide identity, respectively (Fig. 1c). Nearly half of the CrRp3 gene products have the best hits with K1-5 genes, including the DNA and RNA polymerases, DNA ligase and the major capsid protein as well as many hypothetical proteins (Table S3). Accordingly, CrRp3 is considered a new species within the genus *Vectrevirus* in the recently created family *Autographiviridae* [47]. For most phage genera, >95% DNA sequence identity is used by the International Committee on Taxonomy of Viruses (ICTV) as the species demarcation criterion [47]. Gene products unique to CrRp3 include the head-tail connector protein, endolysin, tailspike protein, lyase, minor structural protein and several other proteins with unknown functions (Fig. 1c and Table S1).

### Morphology and genome of phage CrRp10

The second isolated phage, *C. rodentium* phage CrRp10 (formal name: vB_CroM_CrRp10) displays an elongated (prolate) icosahedral head connected to a long tail covered with a discernable sheath (Fig. 2a). These features are characteristic of T4-like myoviruses. Plaques produced by CrRp10 on ICC180 were of <1 mm^2^ (Fig. 2b). This suggests that phage strain has a significant effect on the plaque size, when compared to plaques produced by CrRp3 (Fig. 1b vs Fig. 2b).

**Figure 2.**
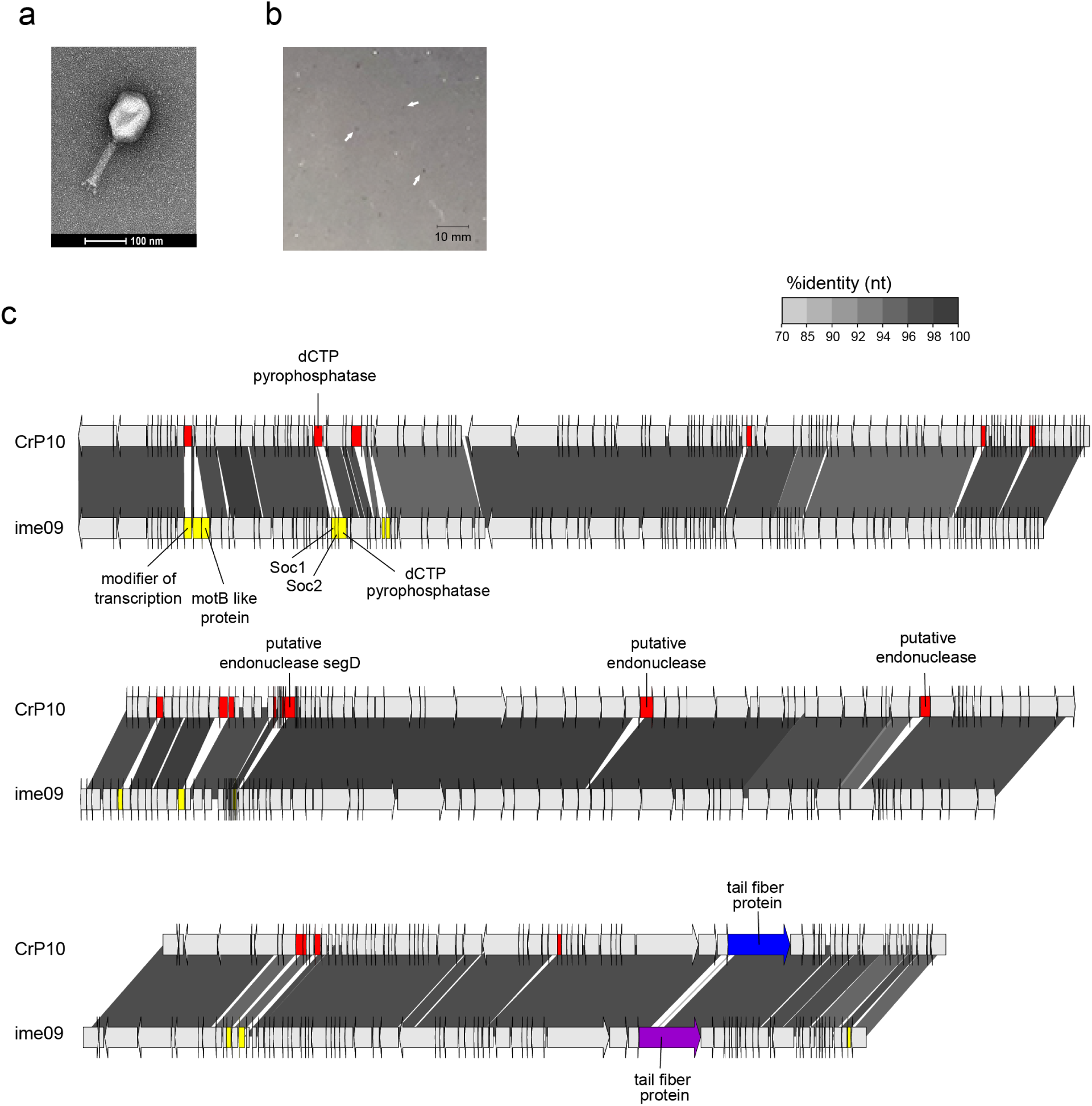
Morphology, genome organization and gene functional comparison of CrRp10. **a)** Electron micrographs of CrRp10 negatively stained uranyl acetate. **b)** Plaque forming units on lawns of *C. rodentium* ICC180. **c)** Alignment of CrRp10 and closed taxonomic relative *E. coli* phage ime09. Genes are colored according to the relationship between CrRp10 and related phage strains, with red labels being exclusive to the CrRp10, yellow labels being exclusive to related phage strain, while blue labels are homologous but highly variable.

Next generation sequencing showed that CrRp10 has a large circularly permuted dsDNA genome of 171.5 kb with 267 CDSs and harbors 10 tRNA genes (Fig. 2c and Table 1). Only 133 (∼50%) of CDS have putative functions (Table S2). Similar to CrRp3, CrRp10 has a GC content of 35.5%, which is 20% lower than that of *C. rodentium.* The genome does not carry sequence homology to lysogeny-associated genes, which suggests CrRp10 is a virulent phage. The genomes also exhibit no sequence homologies to known virulence-associated or antibiotic resistance genes.

At the time of analysis, the closest taxonomic relative of CrRp10 is the *E. coli* phage Ime09, which belongs to the *Tequatrovirus* genus in the subfamily *Tevenvirinae* (Fig. 2c and Table S3). They share significant synteny with 98% nucleotide identity over the complete length, and thus, CrRp10 is considered a new strain. However, little is known about phage Ime09 with details restricted to genomic analysis [48]. CrRp10 has divergence from Ime09 within its tail fiber gene, with 3% amino acid dissimilarity over 80% of the corresponding protein. Another striking feature of the CrRp10 genome is the recombination event, which resulted in the gain of dUTPase with high sequence similarity to that encoded by phage e11/2. Phage e11/2 infects the enterohemorrhagic *E. coli* (EHEC); another A/E pathogen [49]. Other recombination events have added several putative endonucleases with high similarity to homologs in other related Enterobacteriaceae phages (Fig. 2c and Table S2).

### Comparison of Citrobacter phage proteins

Next, we performed amino acid (AA) sequence alignments to compare proteins among phages that infect *C. rodentium* and other *Citrobacter* species (Fig. S3). Surprisingly, CrRp3 has <40% AA homology to either *C. rodentium* phages (CR8 and CR44b) or *C. freundii* phages phiCFP-1 and Sh4), with the exception of homology between a putative lyase (68%) and minor structural protein (75%) of *Citrobacter* phage CR8 (Fig. S3a and Table S3). Several CrRp10 proteins exhibit low AA homology to the *C. rodentium* phage Moon and even less homology with *C. freundii* phage CfP1 (Fig. S3b and Table S3).

### CrRp3 is more likely to find a host, but has smaller burst size

With an excess of host cells, the estimated adsorption rate for phages CrRp3 and CrRp10 are 3.5 ±3.2 ×10^−10^ and 8.52 ±2.8 ×10^−11^ phage^-1^ cell^-1^ mL^-1^ min^-1^, respectively (Table 2). That is, when compared, CrRp3 would ‘find a host’ ∼24 times faster than CrRp10. This also implies these phages have different cell surface binding receptors. In a well-mixed culture, CrRp3 takes approximately 10 min to adsorb, inject its genome, and produce new virions (eclipse period) as well as another 6 min for cell lysis to occur (latent period, which includes phage release) (Table 2). CrRp10 however had eclipse and latent periods of 13 and 18 min, respectively. The shorter replication cycle for CrRp3 correlated with a reduced burst size of 35 phage particles per cell, compared to longer cycling of CrRp10 that produced burst of 70 phage particles (Table 2).

**Table 2.** CrPp3 and CrRp10 replication characteristics on *C. rodentium* strain ICC180.

| Family | CrRp3<br>*Podoviridae* | CrRp10<br>*Myoviridae* |
| --- | --- | --- |
| Adsorption constant<sup>1</sup><br>(phage<sup>-1</sup> cell<sup>-1</sup> ml<sup>-1</sup> min<sup>-1</sup>) | k = 3.50 × 10<sup>-10</sup> (±3.2 × 10<sup>-10</sup>) | k = 8.52 × 10<sup>-11</sup> (±2.8 × 10<sup>-11</sup>) |
| Eclipse period<sup>2</sup> | 10 min | 13 min |
| Latent period<sup>3</sup> | 16 min | 18 min |
| Mean burst size<sup>4</sup> | 34 PFU | 69 PFU |

<sup>1</sup> k = 2.3/Bt \* log(P0/P); t(0) to t(50% bound); ±standard deviation; n=3
<sup>2</sup> The eclipse period represents t(0) to t(intracellular virions prior to cell lysis); n=3
<sup>3</sup> The latent period represents t(0) to t(cell lysis); n=3
<sup>4</sup> n=2

### Hypoxia reduces lysis at low MOIs but not resistance

Both CrRp3 and CrRp10 were able to reduce *C. rodentium* population densities compared to untreated controls under either normoxic or hypoxic culturing conditions (see Methods for details) (Fig. 3a and b, respectively). Under normoxia, CrRp3 took less time to reverse bacterial population growth (∼2.5 h post treatment) and reduced bacterial density below OD limits of detection (LOD) within 3 h (Fig. 3a). For CrRp10, the same took 3.5 and 5.5 h, respectively (Fig. 3b). Of these two phages, CrRp3 appears to be more ‘*potent*’. Under hypoxia, however, CrRp3 at MOIs <0.01 failed to reduce bacterial density below the LOD, whereas CrRp10 could (Fig. 3). Indeed, hypoxia also damped exponential bacterial growth (N0 vs H0) in the absence of phages, but final cell densities were similar after 18 h. In addition, the regrowth of *C. rodentium* occurred between 8-9 h after inoculation with CrRp3. In contrast, no observable bacterial regrowth occurred for CrRp10 after 18 h. Not shown is the significant propagation of both phages in microplate after 18 h compared to the initial inoculums (e.g. 1×10^7^ vs >1×10^9^). These imply that regrowth was due to the emergence of phage-resistance, which was independent of atmospheric conditions.

**Figure 3.**
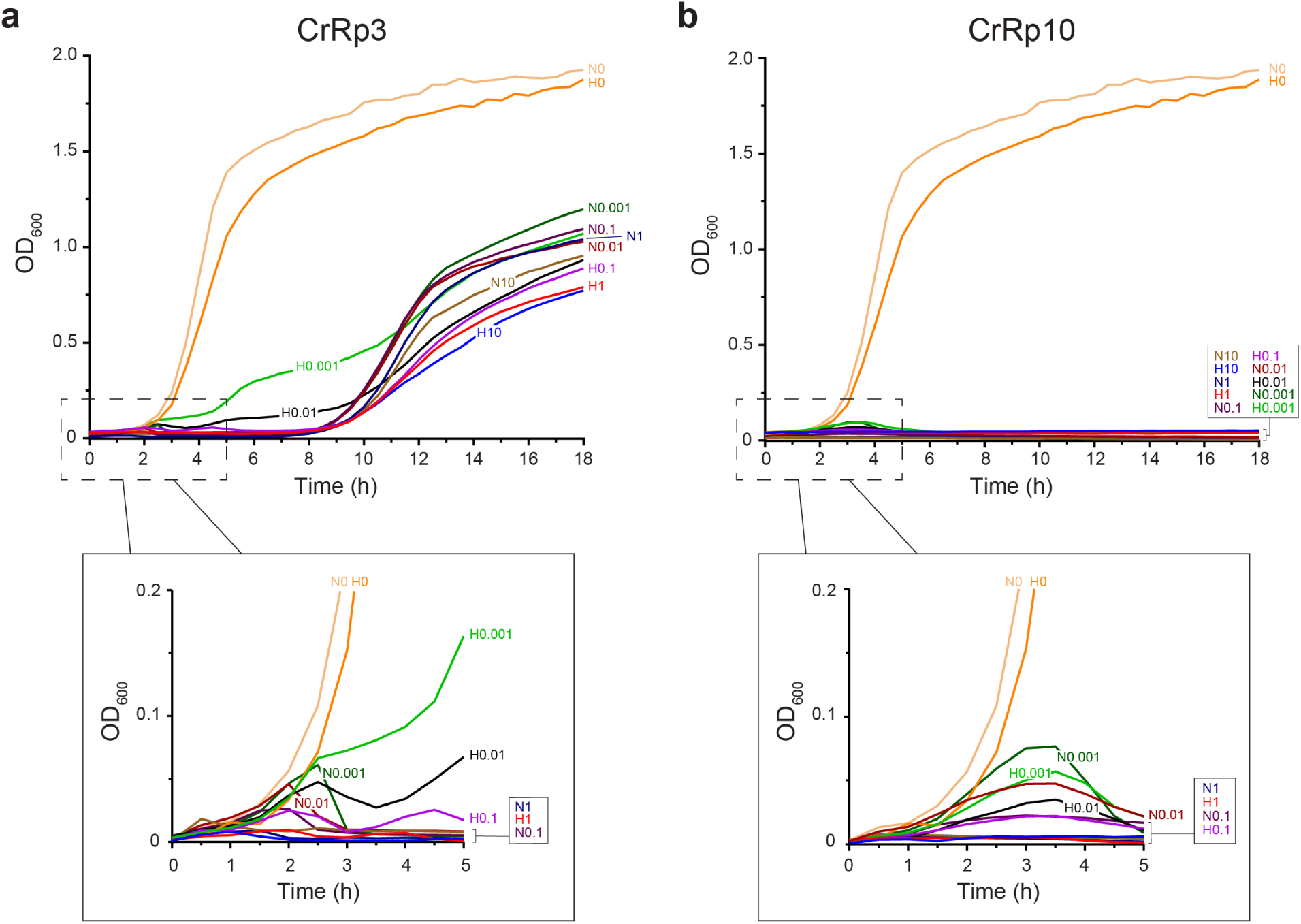
Lysis curves of *C. rodentium* by CrRp3 or CrP10 under normoxic and hypoxic conditions. **a)** Growth curves of *C. rodentium* ICC180 infected with CrRp3 at different MOI (0, 0.001, 0.1, 1 and 10) under either normoxic (N) or hypoxic (H) culturing conditions (see Methods for details). **b**) Growth curves of *C. rodentium* ICC180 infected with CrRp10 at different MOI under either normoxic or hypoxic conditions. Both sub-panels are the first 5 h magnified. n=6

### Host range

Next, we tested the host range of the virulent phages CrRp3 and CrRp10 along with other representative phages against several bacterial strains (Table 3). While, in addition to their isolation strain of *C. rodentium*, CrRp3 could infect the *E. coli* strain K-12, while CrRp10 displays a much broader host range, including K-12 and several pathotypes of *E. coli*, as well as the *E. carotovora* strain CFBP2141. Although the *E. coli* phage LF82_P10 [42] also exhibits a relatively broad host range, it cannot infect *C. rodentium*. Moreover, most of the *E. coli* as well as *P. aeruginosa* and *Serratia marcescens* strains tested were resistant to both CrRp3 and CrRp10.

**Table 3.**
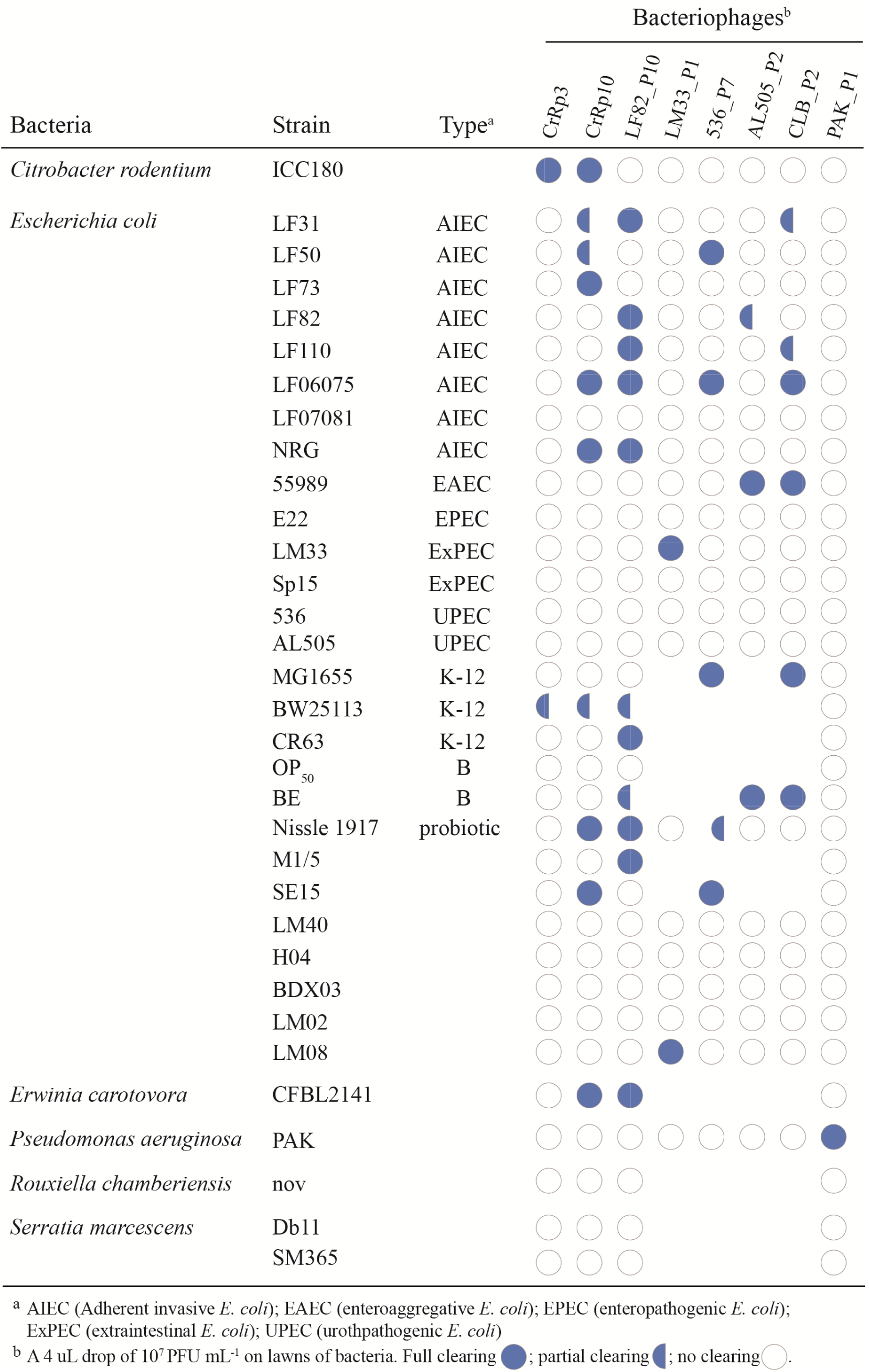
Phage strain host range.

### Phage phylogenetic relationships

To investigate phylogenetic relationships among *Citrobacter* and *Escherichia* podoviruses including CrRp3, we constructed a genome-blast distance phylogeny tree using 37 complete genome nucleotide sequences from GenBank (Fig 5 and Table S4). We found that these phages display heterogeneous clustering of closely related *Autographiviridae* into 3 distinct clades with maximal bootstrap support (Fig. 4a). CrRp3 has closer relationships with podoviruses infecting *Escherichia* (Fig. 4b) than those also infecting *C. rodentium* (CR8 and CR44b) (Fig. 4a; see C1 vs C3). Rather, CR8 and CR44b clusters with some phages that infect the human pathogen *C. freundii* (SH3 and SH4). Other *C. freundii* podoviruses (phiCFP1, SH1, and SH2) cluster in clade 2 (Fig. 4c) and phage CVT2 branches separately. The latter was not unexpected because CVT2 was isolated from the gut of termites with an uncharacterized *Citrobacter* species [50].

**Figure 4.**
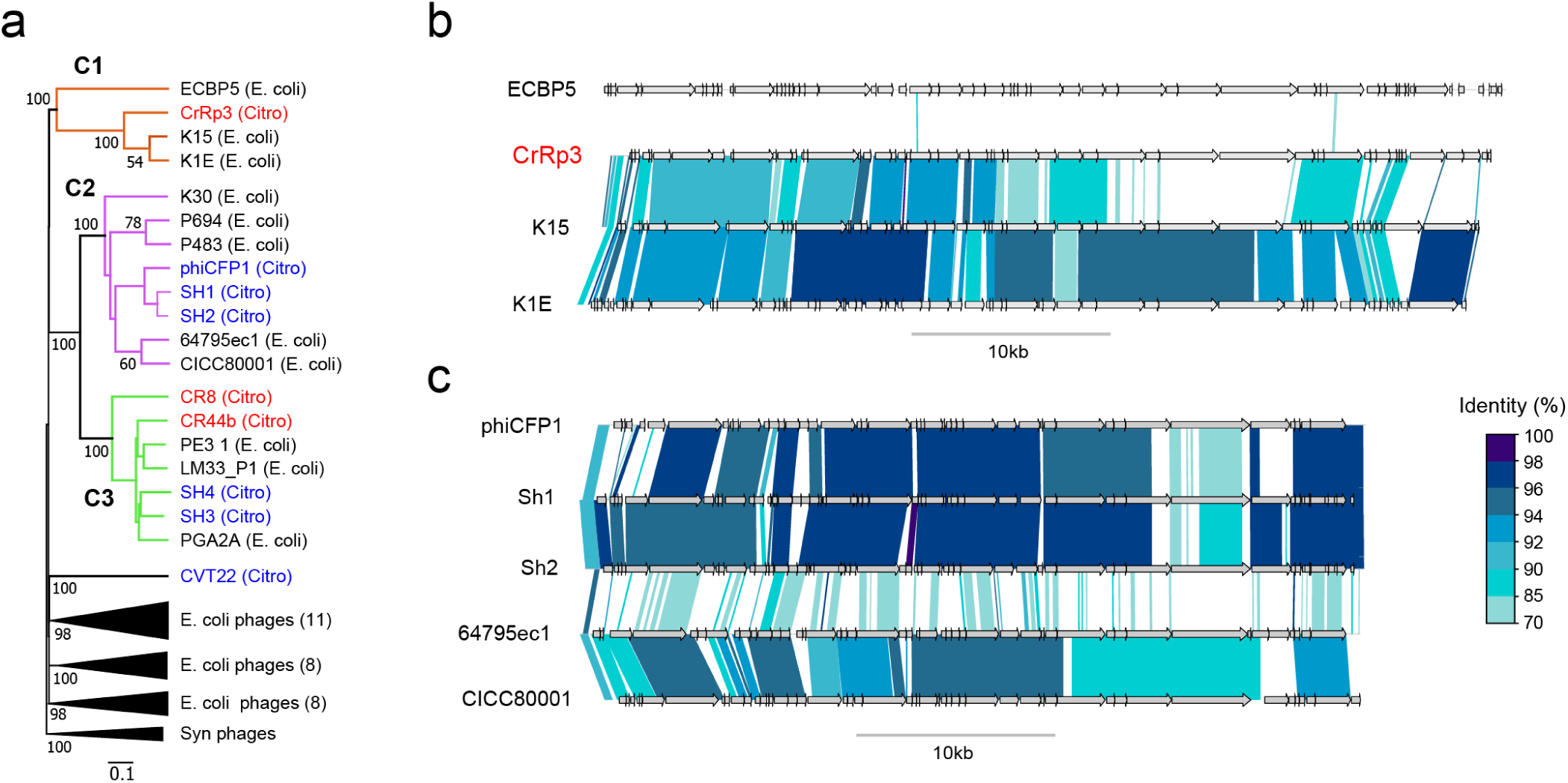
Phylogenetic relationships among *Citrobacter* and *Escherichia* podoviruses. **a)** Tree assembled from complete nucleotide sequences of *C. rodentium* podoviruses (red), other *Citrobacter* spp. podoviruses (blue) and closely related *E. coli* podoviruses are shown. Tree is rooted using *Synechococcus* (Syn) phages and bootstrap values at nodes define percent confidence of 100 replicates. Three phylogenetic clades (C1-3) with strong bootstrap support (>70%) at separating nodes indicated by color. **b)** Detailed nucleotide sequences alignment of members in clade 1 (C1), which includes CrRp3. **c)** Detailed nucleotide sequences alignment of five members in clade 2 (C2), which includes *C. freundii* phages Sh1 and Sh2.

**Figure 5.**
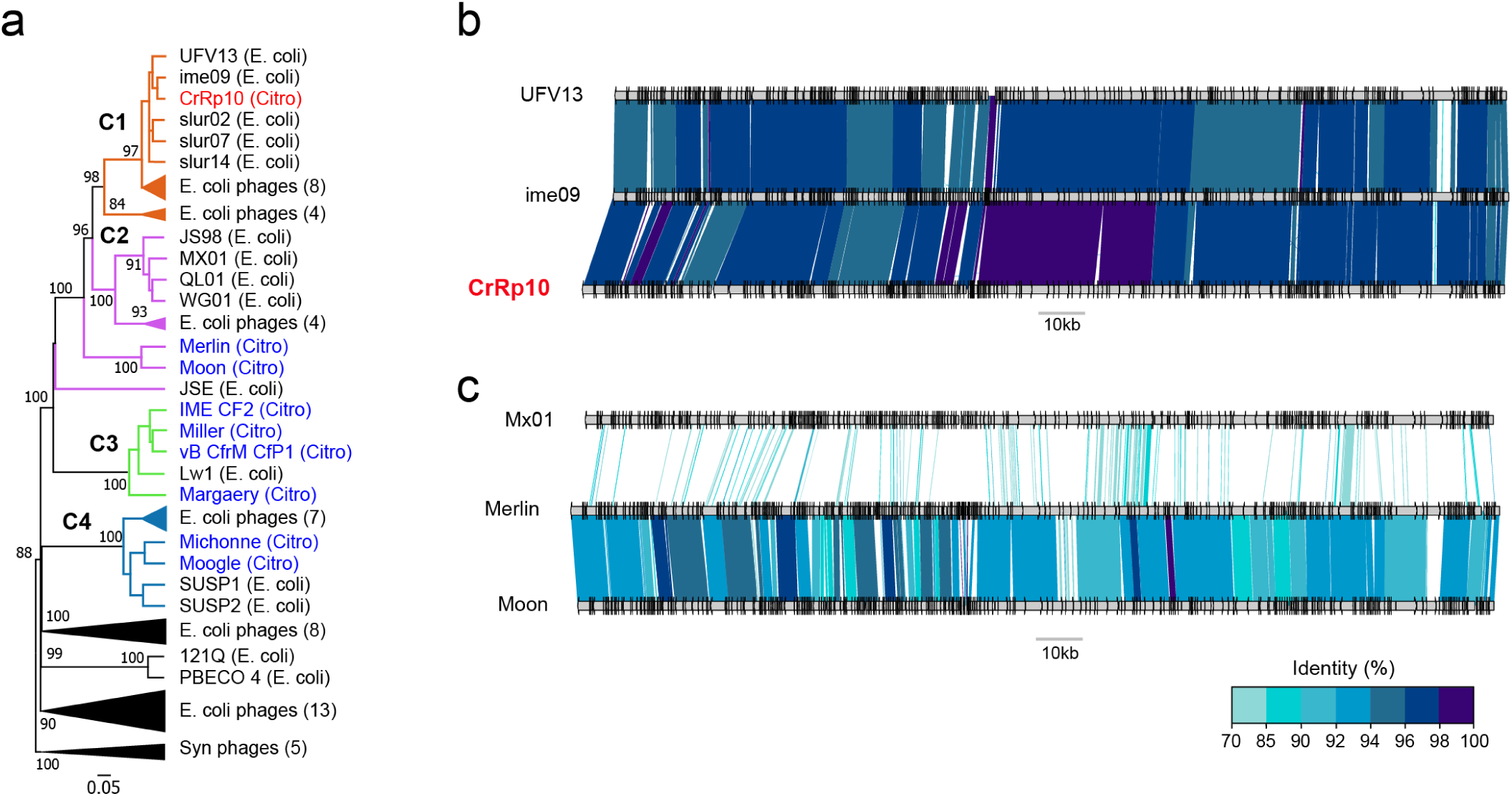
Phylogenetic relationships among *Citrobacter* and *Escherichia* myoviruses. **a)** Tree assembled from complete nucleotide sequences of known *C. rodentium* myoviruses (red), other *Citrobacter* spp. myoviruses (blue) and closely related *E. coli* myoviruses are shown. Tree is rooted using *Synechococcus* (Syn) phages and bootstrap values at nodes define percent confidence of 100 replicates. Four phylogenetic clades (C1-4) with strong bootstrap support (>70%) at separating nodes indicated by color. **b)** Detailed nucleotide sequences alignment of three members in clade 1 (C1), which includes CrRp10. **c)** Detailed nucleotide sequences alignment of three members in clade 2 (C2), which includes *C. freundii* phages Merlin and Moon.

To determine the relationship of CrRp10 to other *Citrobacter* and *Escherichia Myoviridae*, we constructed a genome-blast distance tree using 69 complete genome sequences from GenBank (Fig 5 and Table S4), also showing heterogeneous clustering (Fig. 5a and Supplementary Table S3). CrRp10 clusters with several related *E. coli* myoviruses with high percent nucleotide homology with, for example, phage Ime09 and UFV13 (Fig. 5b). Other distantly related myoviruses that infect *C. freundii* cluster in clades 2-4. Within clade 2 (C2), *C. freundii* phages Merlin and Moon are further distantly related to 8 *Escherichia* phages in C2 (Fig. 5c). In contrast, clade 3 (C3) is composed of almost exclusively *C. freundii* phages (IME CF2, Miller, CfP1, and Margaery).

## DISCUSSION

In this study, we isolated and characterized two new phages, CrRp3 and CrRp10, which infect the mouse-redistricted diarrheagenic pathogen *C. rodentium*. We show that the podovirus CrRp3 is a new species within the genus *Vectrevirus* in the family *Autographiviridae*, while the myovirus CrRp10 is a new strain within the *Tequatrovirus* genus in the family *Myoviridae.* Between CrRp3 and CrRp10, only 48% of their combined gene repertoire have assigned putative functions. Of assignments, we found that neither phage harbor known genes associated with bacterial virulence or antibiotic resistance. The latter is consistent with previous findings that antibiotic resistance genes are rarely carried in phage genomes [51]. In addition, we show that CrRp3 and CrRp10 genomes lack identifiable integrase genes. This implies that both phages carry out strictly lytic replication cycles. Indeed, with more than 50% of genes having unknown function emphasizes the critical need for significant gene functional studies before any certainly that CrRp3 and CrRp10 do not carry harmful genes.

Although, these phages expand our knowledge of viral biodiversity, we found that neither CrRp3 or CrRp10 genomes exhibit nucleotide sequence homology with other *C. rodentium* phages, including CR8 and CR44b from the genus *Caroctavirus* [21]. In addition, CrRp3 and CrRp10 are distantly related to phages that infect *C. freundii*, including members of the genera *Moonvirus* (Merlin, Miller, Moon) [52-54], *Mooglevirus* (Moogle, Michonne, Mordin) [55-57], *Tlsvirus* (Stevie) [58], and *Teetrevirus* (phiCFP-1, SH1, SH2, SH3, SH4, SH5) [59,60]. *C. freundii* can cause a variety of nosocomial acquired extraintestinal human diseases, such as the urinary tract, respiratory tract, and wound infections [61]. Thus, we show that CrRp3 and CrRp10 have independently evolved from closely related *E. coli* phages, presumably because it was advantageous to expand the host range to infect *C. rodentium*, and as a result occupy new niches. The genome of *C. rodentium* exhibits several features typical of recently passing through an evolutionary bottleneck, including several large-scale genomic rearrangements and functional gene loss in the core genomic regions [62-64]. This has led to the hypothesis that *C. rodentium* evolved from human *E. coli*, with the extensive use of mouse models of human *E. coli* infections [64]. Our results strengthen the evidence that *C. rodentium* has relatively recently evolved from *E. coli*. The co-evolution between phages and bacteria could explain why *C. rodentium* phages appear to have emerged from *E. coli* phages, rather than other *Citrobacter* species phages. However, CrRp3 genes have diverged significantly from presumably ancestral genes of *E. coli* phage K1-5, in particular gene products responsible for *C. rodentium* recognition (tail-fibers) and cell lysis (endolysin). Interestingly, phage K1-5 exhibits two tail fiber genes, which allow it to be promiscuous between *E. coli* strains with different capsule compositions [65]. Indeed, CrRp3 endolysin gene has homology to other *Citrobacter* phage endolysin genes. This suggests a mosaic genome structure driven by recombination event from diverse viruses. This is consistent with other autographiviruses that exhibit a high genetic identity and structure and specific RNA polymerase, with the modest differences observed in genes implicated in adaptation to host constraints [66].

Lysis of bacterial cells provides insight into the dynamics between individual phages and their host bacteria. We found that CrRp3 appears to be the more ‘*potent*’ virus of the two. First, it was determined that CrRp3 would be 24x more likely to find a host cell based on adsorption rates compared to than CrRp10. Either this could be due to cell surface receptors for CrRp3 outnumbering receptors for CrRp10 or CrRp3 tail fibers have higher affinity to a shared receptor. Second, in well-mixed cultures, CrRp3 took 2 min less time to complete a single lytic replication cycle than CrRp10 (16 vs 18 min). This correlated to CrRp3 being able to reverse *C. rodentium* exponential growth significantly sooner than by CrRp10 *in vitro* (Fig. 3). This despite CrRp3 exhibiting a 50% reduced progeny burst than CrRp10 (Table 2). This raises the question whether shorter replication cycles are favorable to higher phage progeny production to eliminate bacterial infections.

However, CrRp10 appears more resilient to resistance. Cultures of *C. rodentium* treated with CrRp10 were maintained below the OD limit of detection after 18 h (end of study) even at initial inoculation ratio of 1 phage per 1000 host cells (i.e. MOI = 0.001) (Fig. 3b). In contrast, cultures of *C. rodentium* treated with CrRp3 exhibited regrowth after 9 h at all MOI. Bacteria thwart phage attack through an arsenal of antiviral mechanisms targeting virtually all steps of the phage replication cycle. These include spontaneous chromosomal mutations, the ability to block the entry of phage genetic material (e.g., superinfection exclusion), DNA restriction-modification enzymes, abortive infection and CRISPR-Cas adaptive immunity [67]. El Haddad et al found that 7 of 12 clinical phage therapy studies confirmed resistance had emerged during treatment, including with phage cocktails [68]. For example, *Acinetobacter baumannii* isolated from patient blood cultures exhibited resistance to all eight phages after one week of treatment [69]. However, clinical phage resistance has been under-studied as the root cause of treatment failures. A randomized double-blind clinical trial of a six-phage therapy showed initial reductions in *P. aeruginosa* chronic otitis after treatment, but the infection rebounded in time to pre-treatment levels [70]. Although not tested, the regrowth of bacteria is indicative of emerged resistance to the phages. Thus, selecting phages, like CrRp3, which are resilient to resistance may lead to improvements in phage therapy efficacy.

Moreover, we show that hypoxic conditions that mimic those in animal tissues (2.5–9% O_2_ and 5% CO_2_) [71] caused a notable delay in bacterial population lysis. Enteric pathogens adapt to oxygen limitations by entering into a metabolically altered state [72], which is likely to affect phage-bacteria interactions. For instance, it has been shown that propagation of some phages decreased as the growth rate of the host decreased [73,74]. The effects of hypoxia on phage-bacteria interactions is not completely understood.

Another criterion for selection of therapeutic phages is the spectrum of bacterial species or strains lysed. We found that CrRp3 and CrRp10 exhibit polyvalence in hosts across genera within the *Enterobacteriaceae*. We show that while CrRp3 produced plaques on lawns of *C. rodentium* and the non-pathogenic *E. coli* strain K12, CrRp10 also produce plaques on 8 other *E. coli* strains (pathogenic and non-pathogenic) and the plant pathogen *E. carotovora*. In contrast, most phages are confined to a single host species and often to a subset of strains [43,75]. For example, the *C. rodentium* phage phiCR1 was unable to produce plaques on *E. coli* [19]. Considering the well-documented, collateral effects of broad-spectrum antibiotics, which are notorious for secondary outcomes such as antibiotic-associated diarrhea [76], a narrow phage spectrum may be advantageous during therapy. However, species specificity comes with inherent constraints. By selecting a phage that is limited to a single species or limited number of strains, treatment is likely to be less effective against multispecies or polystrain infections [77].

The overuse and misuse of antibiotics in the treatment of diarrhea has led to an alarming increase of AMR in diarrheagenic bacteria [1-3]. Phage therapy has been shown to be effective at reducing *E. coli* burden in the murine gut with antibiotic pretreatment [5-8]. *C. rodentium* is widely used as an exemplary *in vivo* model system for gastrointestinal bacterial diseases [9,78], but there are no reports using this model in phage therapy development. Prior to animal modeling, careful selection of phages for therapeutic applications was especially important because of the aforementioned drawbacks of potentially carrying unknown toxin genes, the ease of which resistance bacteria can become resistant, the unclear effects of hypoxia of phage-bacteria interactions, and inherent narrow host ranges [12]. It is unclear whether the subtle characteristic differences between CrRp3 and CrRp10 make them ideal for phage therapy. Our hypothesis however is a larger burst size, resistance resilience, and polyvalence of CrRp10 will improve treatment efficacy in mouse models of diarrheal diseases.

## Acknowledgments

The authors are grateful to Drs. Luisa De Sordi and Marta Mansos Lourenço for their advice, acknowledge contributions by Dr. Elena Rensen and grateful to the Ultrastructural BioImaging unit at Institut Pasteur for access to the TEM. This research was supported by an EMBO fellowship (ALTF 1562-2015) and Marie Curie Actions award (LTFCOFUND2013, GA-2013-609409) to CMM and European Respiratory Society grant (RESPIRE2–2015–8416) to DRR.

## SUPPLEMENTARY MATERIALS

(available upon request)

**Supplementary Table S1.** Citrobacter phage CrRp3 annotations.

**Supplementary Table S2.** Citrobacter phage CrRp10 annotations.

**Supplementary Table S3.** Classification of the best hits of phages CrRp3 and CrRp10.

**Supplementary Table S4.** *E. coli* phage complete genomes used in genome-blast distance phylogeny analyses.

**Supplementary Figure S3.**
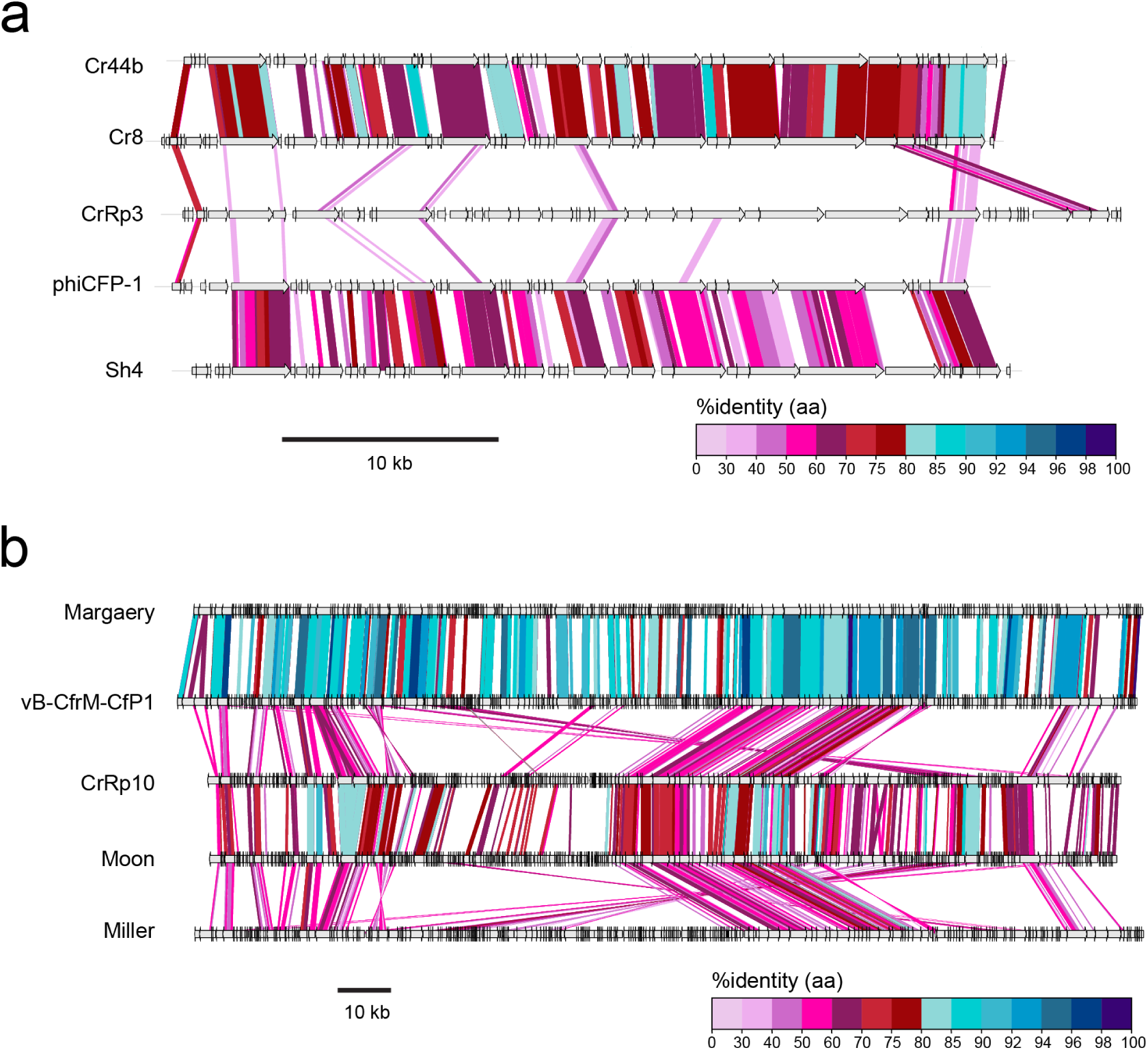
Amino acid sequence alignments to compare the compositions of *Citrobacter* phages a) podoviruses and b) myoviruses.

