## Supplementary figures and images for "Isolation and characterization of bacteriophages that infect *Citrobacter rodentium*, a model pathogen for investigating human intestinal diseases"

### Fig. S1

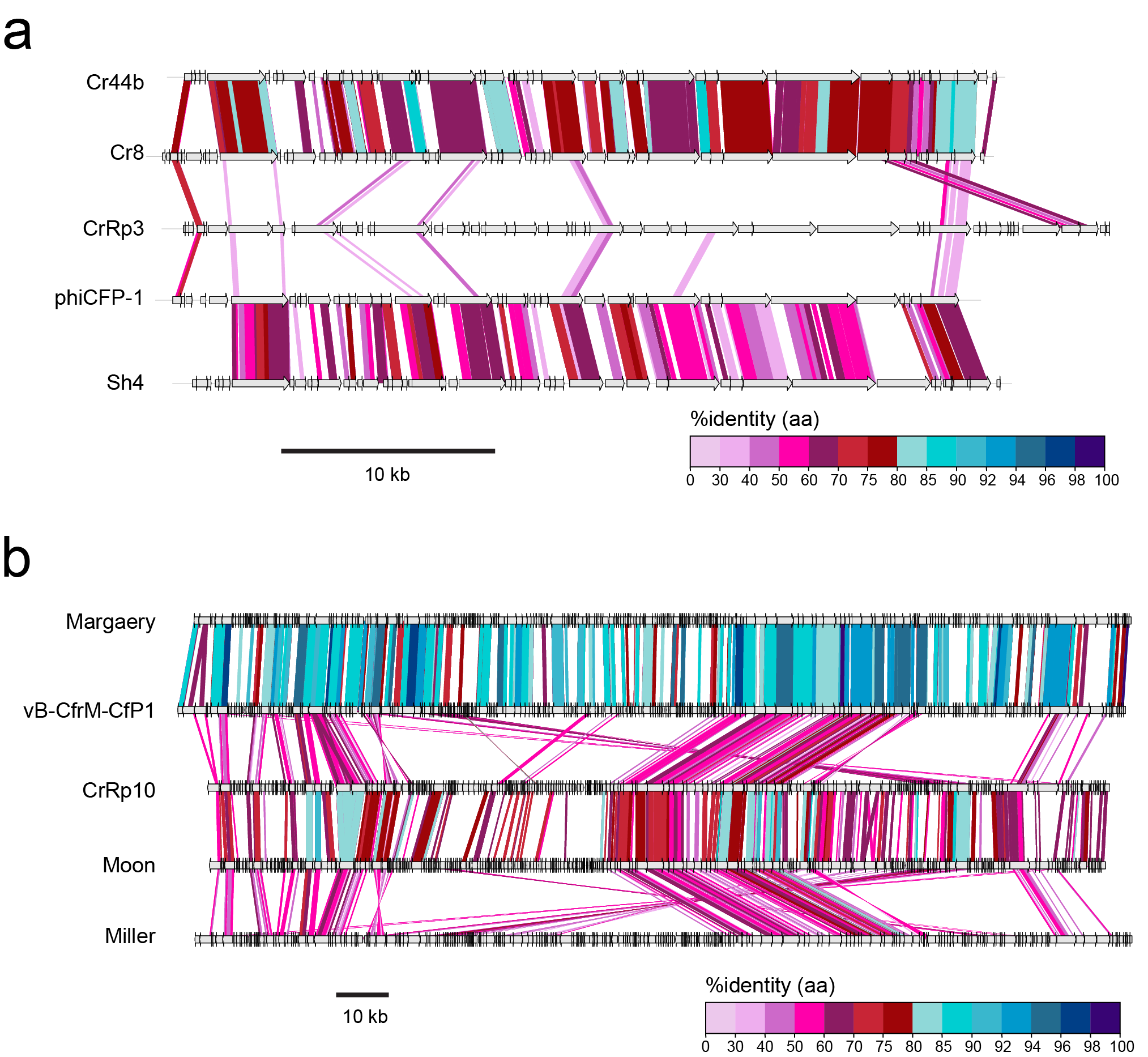
